# The synergetic effect from the combination of different adsorption resins in batch and semi-continuous cultivations of *S. cerevisiae* cell factories to produce acetylated Taxanes precursors of the anticancer drug Taxol

**DOI:** 10.1101/2023.02.06.527354

**Authors:** Jorge H. Santoyo-Garcia, Laura E. Walls, Marissa Valdivia-Cabrera, Koray Malci, Nestor Jonguitud-Borrego, Karen J. Halliday, Leonardo Rios-Solis

## Abstract

*In situ* product recovery is an efficient way to intensify bioprocesses as it can perform adsorption of the desired natural products in the cultivation. However, it is common to use only one adsorbent (liquid or solid) to perform the product recovery. For this study, the use of an *in situ* product recovery method with three combined commercial resins (HP-20, XAD7HP and HP-2MG) with different chemical properties was performed. A new yeast strain of *Saccharomyces cerevisiae* was engineered using CRISPR Cas9 (strain *EJ2*) to deliver heterologous expression of oxygenated acetylated taxanes that are precursors of the anticancer drug Taxol ® (paclitaxel). Microscale cultivations using a definitive screening design (DSD) were set to get the best resin combinations and concentrations to retrieve high taxane titers. Once the best resin treatment was selected by the DSD, semi-continuous cultivation in high throughput microscale was performed to increase the total taxanes yield up to 783 ± 33 mg/L. The best T5α-yl Acetate yield obtained was up to 95 ± 4 mg/L, the highest titer of this compound ever reported by a heterologous expression. It was also observed that by using a combination of the resins in the cultivation, 8 additional uncharacterized taxanes were found in the gas chromatograms compared to the dodecane overlay method. Lastly, the cell-waste reactive oxygen species concentrations from the yeast were 1.5-fold lower in the resin’s treatment compared to the control with no adsorbent aid. The possible future implications of this method could be critical for bioprocess intensification, allowing the transition to a semi-continuous flow bioprocess. Further, this new methodology broadens the use of different organisms for natural product synthesis/discovery benefiting from clear bioprocess intensification advantages.

## 1. Introduction

Microbial cell factories are becoming an important route for the sustainable production of different natural products and chemicals of different commercial (therapeutics, food additives, cosmetics or biofuels) or academic relevance (Malcı et al., 2020; Walls, Otoupal, et al., 2022; Wong et al., 2017, 2018). To date, a majority of studies involving an integrated process of product synthesis and recovery have not been performed at a lab scale, which has contributed to the slow translation of microbial cell factories to fulfil industry demands (Walls, Martinez, & Rios-Solis, 2022; Walls, Martinez, del Rio Chanona, et al., 2022). Different bioprocess strategies have been used to enhance the recovery of desired products, for instance, using two or more phase partitions, foam phase recovery or *in situ* product recovery (ISPR). ISPR is a common integration bioprocess strategy that incorporates a step that sequesters microbe products released to the media, making the production and extraction a single-step method (De Brabander et al., 2021). Many studies have developed an ISPR using liquid solvents, such as dodecane (Ajikumar et al., 2010), polypropylene glycol 4000 (Nicolaï et al., 2021), monoterpenes (Brennan et al., 2012) or ionic liquids (Zhong et al., 2012). The use of a liquid extractant is very practical if the solvent has an affinity to the molecule of interest (Najmi et al., 2018). However, this method has some limitations, such as the liquid extractant must be non-toxic to microbial cell factories, and the product recovery from these solvents can be problematic as they can be hard to evaporate such as dodecane, while not being environmentally friendly (Santoyo-Garcia et al., 2022; Walls, Martinez, del Rio Chanona, et al., 2022).

In addition, solid extractants for ISPR have also been widely used, where common materials used to achieve the recovery of products involve: activated charcoal, silica gel and adsorbent porous resins among others (Lee et al., 2003; Othman et al., 2018). An important advantage of using a solid extractant is the easiness to be separated from the media by centrifugation or filtration steps as well as the easy recovery of the desired product from the solid phase (Aguilar et al., 2019). Various studies have assessed the efficacy of single resin beads via *in situ* cultivation. Phillips et al., 2013 showcased the use of several resins (XAD-7, XAD-8, XAD-1180, HP-20 and HP-21 among others) *in situ* with different microorganisms (*Streptomyces spectabilis, Penicillium chrysogenum, Salinispora tropica, Sorangium cellulosum, Actinoplanes teicomyceticus* among others) to get an ample range of natural products such as anticancer drugs (Epothilones, Leinamycin, Dynemicin A among others), antibiotics (Streptovaricin, Pristinamycin, Brotrallin, etc.), antioxidants (anthocyanin, stilbenes, among others) and many more (Luo et al., 2014; Phillips et al., 2013; Shin & Kim, 2016). One of the most studied resins for ISPR cultivation is HP-20, which was shown to increase the yields of anticancer drugs such as thailandepsin from less than 10 mg/L in *Burkholderia thailandensis* to 237 mg/L (Liu et al., 2012).

Another advantage of ISPR, and more specifically the use of solid extractants, is the potential capacity of cell-waste removal in addition to product recovery, which could be crucial to optimize the bioprocesses (Ojovan et al., 2011). The most common cell-waste generators are ethanol, ketones, ammonia, and lactate among others (De Brabander et al., 2021), as they have been linked to poor cell growth or product inhibition if their concentration surpasses any established tolerance level (Wang et al., 2011). Another undesired effect of cell waste is the unintended activation of stress genes that affect cell homeostasis, oxidative stress and reactive oxygen species (ROS) (Rodacka, 2016). As previously mentioned, ISPR could alleviate the waste load produced during the cultivation by selectively adsorbing it into the resin beads surface or the solvent extractant. While there are clear advantages to this strategy, the value of using a combination of solid extractants has not been sufficiently investigated. By contrast, the combination of solvents has been more commonly used in the extraction step by coalescing solvents to maximize the recovery of the targeted molecule (González-Peñas et al., 2014; Park et al., 2007). In previous studies, the blend of solid extractants had been used primarily for the removal of hard-chemical industrial waste from water or for capturing CO_2_, therefore, this approach has been more studied in the environmental field rather than for industrial bioprocess intensification (Brennan et al., 2012; Marin & Stanculescu, 2022; Mortezaeikia et al., 2015).

Finally, another key advantage of using solid extractive methods is the natural transition towards continuous or semi-continuous fermentation systems with ISPR. Maintaining the solid extractants during semi-continuous cultivation, the evaporation of the extractant could be minimized, which has been reported as a common drawback in the dodecane overlay method (Nowrouzi et al., 2020; Walls et al., 2020, 2021).

This study, therefore, set out to assess different resins with polar, non-polar and aliphatic adsorption ranges for ISPR to enhance the cell-waste and product removal. As there are few studies focusing on the optimization of combining solid extractants with different polarities in cultivations using microorganisms, this work aims to follow a Design of Experiment (DoE) approach to test the effect of three commercial adsorbent resins simultaneously on the total taxanes recovery, further allowing the adsorption for uncharacterized or uncharacterized secondary metabolites. Further, this study seeks to maximize total taxanes titers in a high throughput system by using semi-continuous cultivation on a microplate scale. Three resin beads were used simultaneously for engineered *S. cerevisiae* fermentations with ISPR, these include Diaion © HP-2MG (polar), Diaion © HP-20 (non-polar) and Amberlite © XAD7HP (aliphatic) in a range of concentrations and combinations to find the effect on cell growth and total taxane recovery. HP-2MG is an adsorbent resin that is made mainly of methacrylic ester copolymer giving it a higher polarity, it counts with an average surface area of 500 m^2^/g, an average particle size between 500 and 600 μm, a pore size of 170 Å (Unsal et al., 2012). HP-20 resin beads are conformed by polystyrene and divinylbenzene with no oxygen in their structure giving them non-polarity, they have a mean surface area of 720 m^2^/g, a particle size between 400 −1000 μm, a pore size of 290 Å (Aguilar et al., 2019). Finally, XAD7HP resin is primarily made of acrylic ester giving it a medium polarity between the previous resins, it has an average surface area of 450 m^2^/g, an average particle size of 560 and 710 μm, a pore size of around 90 Å (Marin & Stanculescu, 2022). A schematic description of the reaction pathway and methodology followed in this study to perform the ISPR can be seen in Figure 1.

**Figure 1.**
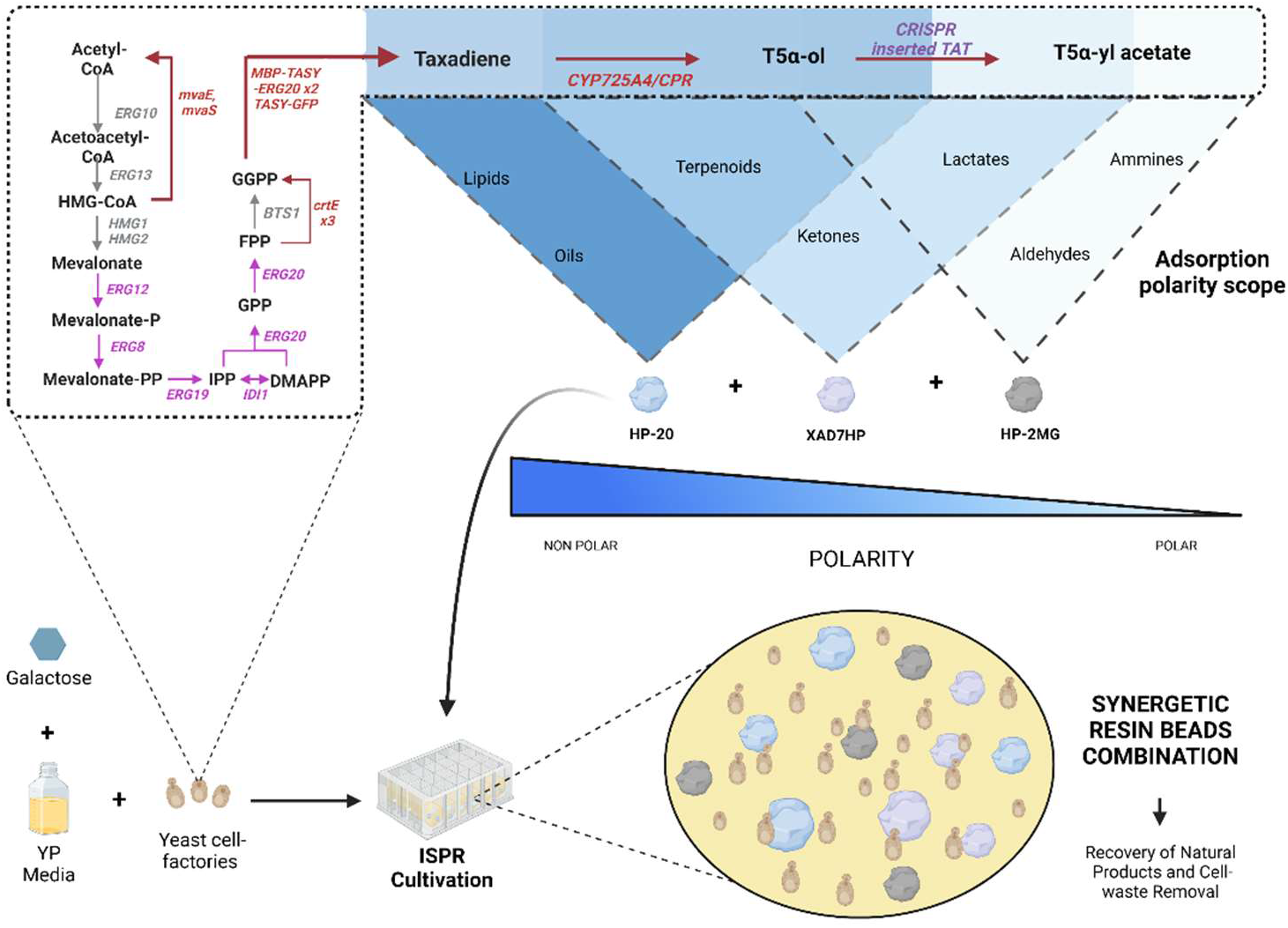
Different adsorbent porous resin beads were used simultaneously for an engineered *S. cerevisiae* fermentation to produce and recover taxanes. By combining the adsorption capacities of the three resin beads (Diaion ©HP-20, Amberlite © XAD7HP and Diaion © HP-2MG), it was intended to increment biomass accumulation and the total taxanes production by helping the cell remove cultivation wastes while improving product extraction.

This research was focused on constructing an engineered *S. cerevisiae* strain *EJ2* by modifying the previously built strain *LRS6* (Walls et al., 2021) that was able to produce the oxygenated Taxadiene-5α-ol taxane. An extra copy of the gene of Taxadiene-5α-ol O-acetyltransferase was integrated into the genome using CRISPR-Cas9 to produce Taxaxadiene-5α-yl acetate (Figure 1). Finally, after optimizing the bioprocess using DoE in microplate batch cultivations, the best concentration and combination of resin beads was selected for further semi-continuous microplate cultivations to optimize the methodology according to total taxanes and oxygenated acetylated taxanes recovered.

## 2. Materials and Methods

### 2.1 Yeast Strain and Media

The *S. cerevisiae* strains used in this study were a CEN.PK2-1C-originated yeast strain (EUROSCARF, Germany), and a CEN.PK2-1C yeast strain previously engineered by (Walls et al., 2021) labeled as *LRS6* with the following genotype: MATa, leu2-3, 112::HIS3MX6-GAL1p-ERG19/GAL10p-ERG8; ura3-52::URA3-GAL1p-MvaSA110G/GAL10p-MvaE; his3Δ1::hphMX4-GAL1p-ERG12/GAL10p-IDI1; trp1-289::TRP1_GAL1p-CrtE (X. dendrorhous)/GAL10p ERG20; YPRCdelta15::NatMX-GAL1p-CrtE/GAL10p-CrtE; ARS1014::GAL1p-TASY-GFP; ARS1622b::GAL1p-MBP-TASY-ERG20; ARS1114a::TDH3p-MBP-TASY-ERG20; ARS511b::GAL1p-T5αOH/GAL3-CPR. Using the CRISPR Cas9 methodologies described by (Malci et al., 2022) Malci et al. (2022), the strain *EJ2* was created with the following genotype (*LRS6*, RKC3::GAL1-TAT).

All cultivation reagents were sourced from Fisher Scientific UK at the highest available purity unless otherwise stated. The media used for the yeast cultivations was yeast extract-peptone (YP, yeast extract 1 % (w/v), peptone 2 % (w/v) supplemented with 2 % (w/v) galactose (YPG) or 2 % (w/v) glucose (YPD) unless otherwise stated..

### 2.2 Batch cultivation at microscale

Microscale cultivation screening was performed using 24-well 10 mL deep well plates (Axygen, USA) with a working volume of 2 mL. Inoculum cultures were prepared by transferring a single colony of the relevant *S. cerevisiae* strain to 5 mL of YPD media and incubating in an orbital shaker incubator (Innova 42, Eppendorf, UK) at 30 °C and 250 rpm overnight. An aliquot of this culture was then diluted with YPG media to give a 2 mL culture with an initial OD_600_ = 1. Adsorbent resin beads Diaion® HP-20 (Merck, Germany), Amberlite © XAD7HP (Merck, Germany) and Diaion © HP-2MG (SUPELCO, UK) were added in different combination sets and concentrations (Section 2.8). Before use, the resin beads were autoclaved at 121 °C for 15 minutes. The resulting culture microplate was incubated at 30 °C for 72 hours at 350 rpm in a Thermomixer C (Eppendorf, Germany) and covered with an adhesive gas permeable membrane (Thermo Fisher Scientific, UK). At the end of the cultivation, the contents of each well were centrifuged at 4500 rpm for 10 minutes, before the supernatant was separated from the pellet (biomass + beads). Biomass accumulation was monitored as optical density at 600nm via offline sampling using a Nanodrop 2000c spectrophotometer (Thermo Fisher Scientific, UK) with a dilution factor of 20 μL of sample in 980 μL of distilled water. To ensure the presence of beads did not inflate optical density readings, controls containing culture media and resin beads in the absence of cells were run in parallel. As the OD_600_ readings of these controls were not different from the blank, a correction was not deemed necessary in the culture readings. Control YPG cultivations with no extraction aid were also performed at the same microplate in different wells. Dodecane overlay cultivations were performed under the same conditions as controls with no extraction aid but with the addition of 20% of the cultivation volume of sterile dodecane at 99% purity (Acros Organics, UK). All microplate cultivations were made in triplicates to increase the statistical robustness of the results, with standard deviation represented as error bars.

### 2.3 Biomass and resin beads partition for taxane quantification

The partition of resins and biomass was made after the batch cultivations in ISPR treatments. Partition was performed using a 200 μL micropipette where the yellow tip was placed as close to the microplate bottom as possible. Once the tip was in direct contact with the plastic wall of the microplate, media and biomass removal was conducted. The space generated between the tip and microplate and the top of the micropipette was small enough to restrict the passing of any resin beads with a diameter bigger than the 400 μm of the resins used in this study. Each separated phase (biomass and resins) was then put into the extraction of taxanes protocol detailed in section 2.4.

### 2.4 Extraction of taxanes

Extraction of taxanes was performed after the cultivation process using only the solid phase (biomass + beads), which was recovered and dried under nitrogen gas at room temperature before adding acetone (99.8%, Thermo Fisher Scientific, UK) at a 1:1 ratio of the cultivation volume and incubating the resulting mixture at 30 °C and 350 rpm for four hours. Following the extraction incubation, the mixture was centrifuged at 4500 rpm for 10 minutes and the organic phase was recovered for taxadiene analysis via Gas Chromatogram -Mass Spectrometry (GC-MS). Taxanes from control runs were extracted from the solid phase (biomass only) at the end of the cultivation using acetone at a 1:1 ratio. The resulting mixture was incubated at 30 °C and 350 rpm for four hours and sent for GC-MS analysis.

### 2.5 Ammonia quantification

After the cultivation process and centrifugation, the supernatant was separated for ammonia quantification. Nessler’s Reagent (Potassium tetraiodomercurate) was used to measure the ammonia ions in the culture media. An aliquot of 10 μL of the supernatant sample was transferred to a 2 mL microcentrifuge tube, and then 200 μL of Nessler’s Reagent (Merck, Germany) was added to the same tube along with 100 μL of Rochelle salt (KNaC_4_H_4_O_6_ × 4H_2_O) to minimize other ion interferences (Zhao et al., 2019). Finally, 490 μL of deionized water was added to dilute the sample into a readable range of the spectrophotometer. The tube was mixed using a vortex for 30 seconds and left at room temperature (25 °C) for 25 minutes. At the same time, a blank preparation was made using the same reagents but substituting the same sample volume with deionized water. Samples were transferred to 1 mL cuvettes and run through a Nanodrop 2000c spectrophotometer (Thermo Fisher Scientific, UK) using a UV-VIS at 420 nm. Obtained values were then added to the previously made calibration curve to calculate the final concentration of ammonia in the culture media. A calibration curve was made using the same protocol with ammonia chloride 99.998% (Merck, Germany) diluted in different calibration points as a standard solution. The calibration curve equation values and calibration points can be seen in Figure S9 in the supplementary material.

### 2.6 Reactive oxygen species (ROS) quantification

ROS quantification was made at different cultivation points for both treatments and controls (at 24, 48 and 72 hrs.). The intracellular fluorescence method (Abhishek & Elah, 2021) was used with some modifications. At the specific cultivation times, a homogenous aliquot of 100 μL was separated from the rest of the cultivation media. Then, 4 μL of 2’, 7’-dichlorodihydrofluorescein diacetate (H_2_-DCFDA) solution (Invitrogen, USA) was added from the stock solution (5 mg/ml) and incubated at 30 °C for one hour. The mixed solution was centrifuged at 4500 rpm for 5 minutes to keep the pellet which was resuspended and washed twice with 500 μL of PBS 1X (Merck, Germany) to finally add 200 μL of PBS. This solution was then transferred to a dark 96 well plate (Greiner Bio-one, Germany) for fluorescence reading by excitation at 484 ± 10 nm and emission at 518 ± 10 nm wavelengths in a microplate reader (FLUOstar Omega, BMG Labtech).

### 2.7 Semi-continuous microplate cultivation

Semi-continuous cultivations were performed at microscale, with variables such as: working volume, media constitution, initial OD and resin beads used were maintained, as described in section 2.2. After batch cultivations, the best resin combination ratio was selected (0.5, 1 and 1.5% (w/v) of HP-20, XAD7HP and AH-2MG respectively) and maintained for further experiments. Different maximum concentrations of resins were used (3%, 6% and 12% (w/v)) to test the resins adsorption saturation of the total taxanes. Before reaching the stationary phase (∼ 48 hours), 75% of the working volume was carefully removed using 200 μl tips without removing any resin. Afterwards, the same removed volume was replenished with fresh YPG media. This step was performed three times before the cultivation was left to run for a final 72 hours, resulting in a 9-day total cultivation process. This procedure is further detailed in Figure S1 and Table S1 from the supplementary material. Sacrifice sampling was used in triplicates to determine the total taxanes adsorption kinetics. At the end of the cultivation, extraction was performed as described in section 2.3 for taxanes analysis in the GC-MS.

### 2.8 Analytical methods

Taxanes identification and quantification was achieved via GC-MS. A 1 μL sample of each organic solvent extract was injected into a TRACE™ 1300 Gas Chromatograph (Thermo Fisher Scientific, UK) coupled to an ISQ LT single quadrupole mass spectrometer (Thermo Fisher Scientific, UK). Chromatographic separation was achieved using a Trace Gold TG-SQC gas chromatography column (Thermo Fisher Scientific, UK) using a previously described method (Walls et al., 2021). To identify and quantify total taxane production by the *S. cerevisiae* strain *EJ2*, pure standards of taxadiene, kindly supplied by the Baran Lab (The Scripps Research Institute, California, USA), and GGOH, obtained from Sigma Aldrich (Gillingham, UK), were used. Characteristic gas chromatograms from different methods and taxane mass spectra can be found in the supplementary material (Figure S2 to S8).

### 2.9 Design of experiments; Definitive Screening Design (DSD) for microplate cultivations and statistical analysis

To test the effects of different factors in microplate cultivations, JMP data analysis software (SAS) was employed for both design of experiments and statistical modelling. A three-level definitive screening design (DSD) capable of screening second-order effects was used. The numerical values of total taxanes obtained from the GC-MS TRACE™ 1300 Gas Chromatograph (Thermo Fisher Scientific, UK) were used as a response in the designs. The order of the conditions was randomized and a minimum of 14 treatments were tested, and the three adsorbent resin concentrations were set as continuous factors. Forward stepwise regression was used (using a α value of 0.05) to make the construct model effects. The ‘response surface’ at the Macros option was selected to test all possible factor’s interactions, with the minimum Bayesian information criterion (BIC) as a stopping rule for stepwise regression control. For optimization, the desirability score was maximized, and the parameters suggested by the models were used to find the highest taxane titers in semi-continuous cultivation. No resin or extraction aid *in situ* in the cultivation of the *S. cerevisiae* strain *EJ2* was used as control. Experiments were conducted in triplicate; error bars indicate the standard deviations within the samples. Detailed information about the DSD can be seen in Table 1.

**Table 1.**
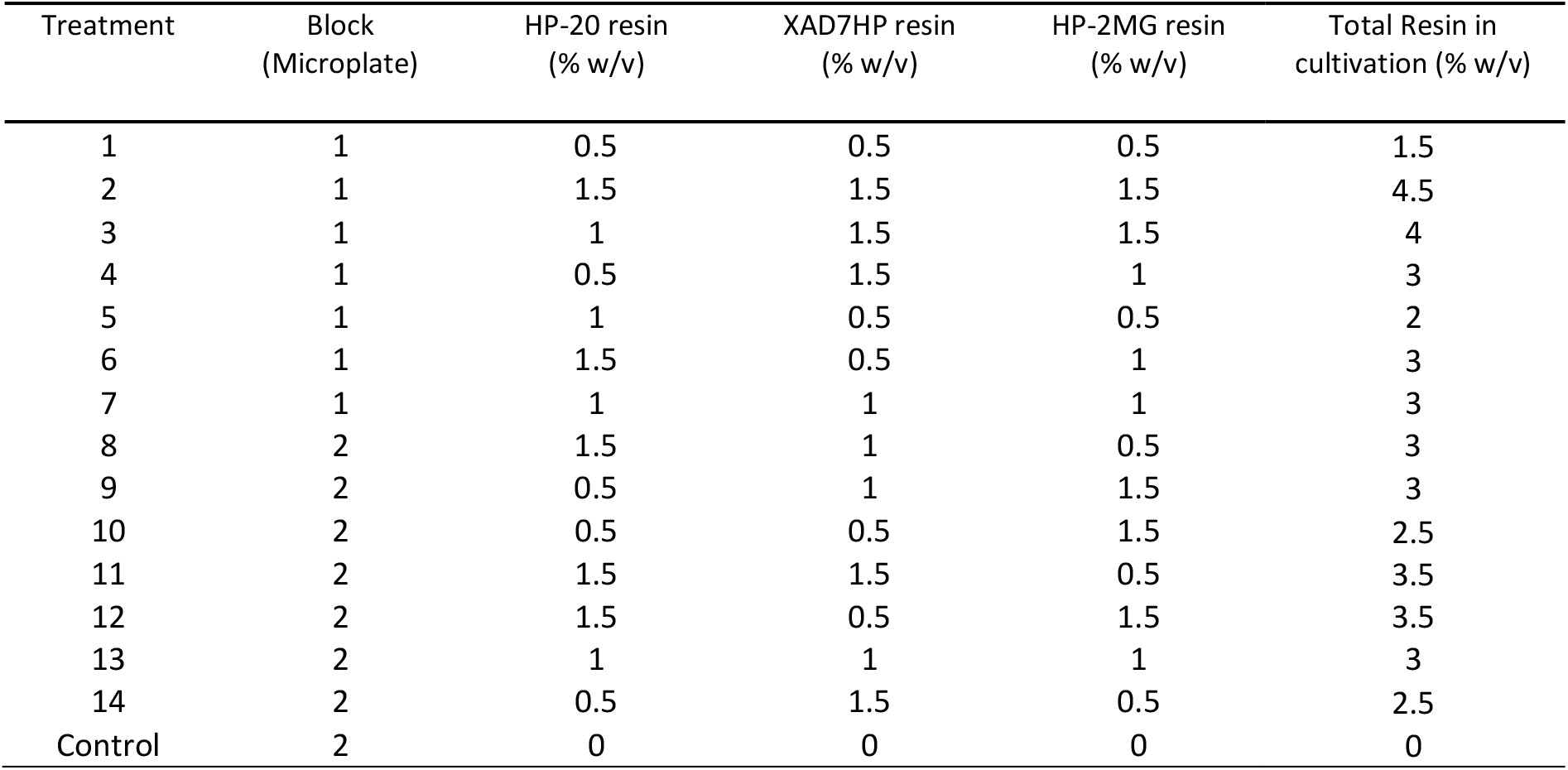
Experimental Design Factors and levels. These results were obtained from the Definitive Screening Design (DSD).

Statistical analyses were performed using GraphPad Prism 8.0 software. One-way analysis of variance (ANOVA) was used to determine whether combined resin bead treatments yielded a significant impact on biomass accumulation and cell-waste removal compared with controls that lacked any extraction aid. A Dunnett’s multiple comparison test was used to compare each treatment to the control. One-way ANOVA was also used to determine whether the quantity of resins was significant in the total taxane titers in the semi-continuous runs. A Tukey’s honest significant difference test was subsequently employed to compare the total resins concentrations of 3, 6 and 12% (w/v) of treatment 9 (Table 1) in the batch and semi-continuous experiments. The null hypothesis considered that there was no significant difference between the treatments, hence if *p* ≤ 0.05 the null hypothesis was rejected.

## 3. Results and discussion

### 3.1 Design of experiments to optimize the resin beads ratio and concentration based in the cell growth and total taxane production

The *in situ* extraction material used in this study has been used separately in previous studies for product recovery enhancement (Halka & Wichmann, 2018; Lee et al., 2003; Qi et al., 2014). However, the use of these materials must be well studied as they can negatively affect cell growth and reduce the efficiency target molecule recovery (Othman et al., 2018). Therefore, to optimize this, a set of experiments were performed in the presence of different resin beads to investigate which combination resulted in the highest biomass accumulation after 72 hrs. Controls in these experiments were carried out with no adsorbent during the cultivation process. To thoroughly investigate the effect of combining the three resin beads, a Definitive Screen Design was implemented using three factors (resin types) at different concentrations, taking into account cultivations of around 3% (w/v) have been shown as the optimal resin bead concentrations from previous work (Santoyo-Garcia et al., 2022). The results on the effect of cell growth and total taxane production are summarized in Figure 2.

**Figure 2.**
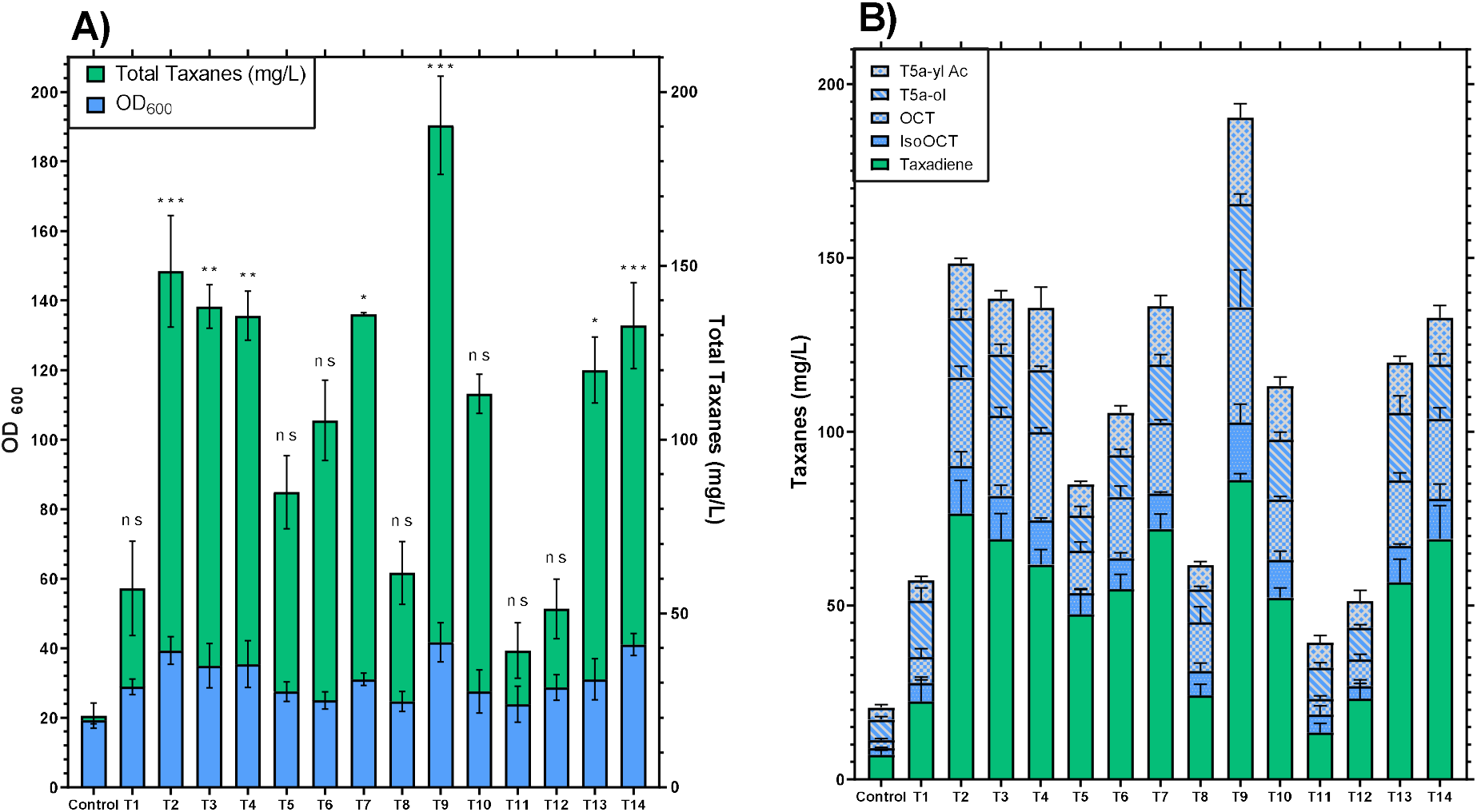
*S. Cerevisiae* strain *EJ2* growth and total taxane titers according to DSD microplate cultivations. A) Cultivations were performed using 2 mL working volume in microplates, YPG and different sets of resin beads combinations which are detailed in Table 1. All cultivations were performed in microscale at 30 °C and 350 rpm shaking using HP-20, XAD7HP and HP-2MG as solid extractants. Blue bars represent OD_600_ at the end of cultivation and green bars represent the total taxanes retrieved. B) Breakdown of the total taxanes retrieved in all treatments after 72 hrs. of cultivation. Blue bars represent the oxygenated taxanes and green bars the non-oxygenated taxanes. Bar values represent mean ± standard deviation (n = 3). Statistical significance for biomass accumulation was assessed by Dunnett’s test versus their corresponding controls. *P <0.0332; **P <0.0021; **P <0.001 and n.s.= not significant P>0.1234.

The different combinations of the resin beads HP-20, HP-2MG and XAD7HP were found to increase the total biomass concentration in *S. cerevisiae* strain *EJ2*, compared to the control, in almost all the conditions of the DoE (Figure 2A). The resin bead combinations that had a higher biomass accumulation at the end of cultivation were treatments 2, 9 and 14 (p < 0.001), showing an OD_600_ of 39 ± 3, 42 ± 4.5 and 40 ± 3 respectively, compared to the OD_600_ of 19 ± 1 in the control cultivation without beads (Figure 2A). When single resins were used, OD_600_ values were 31 ± 1, 34

± 0.5 and 34.5 ± 1 for the resins HP-20, XAD7HP and HP-2MG respectively (Figure S10 of supplementary material), meaning the resin combination treatment would still have higher biomass accumulation at treatments 2, 9 and 14. These results indicated that a favorable synergetic effect is been performed by the combination of resin beads in relation to cell growth. This positive effect could be due to the removal of toxic waste by selected beads during the cultivation of the yeast. Also, the adsorption range of distinct possible toxins is potentially heavily increased by using different resin beads, having a potentially positive effect on cell growth. These results showed that the use of all resin types was beneficial for cell growth and natural products biosynthesis (Figure 2A), supporting the results reported by Aguilar et al. (2019), who tested different adsorbent resins *in situ* for the recovery of Sesquiterpene (+)-Zizaene in engineered *E. coli*. All resins used by these authors individually improved biomass accumulation at the end of the fermentation by 1.35-fold compared to a control with no solid extractants (Aguilar et al., 2019). The use of solid adsorbents at some point during the cultivation resulted in favorable biomass accumulation, although the advantage of combining adsorbents during the cultivation had not been seen in recent studies. Apart from increasing cell growth yield, modifying the combination of beads could also be advantageous depending on the target molecule for recovery. An example of this is that by using the resin combination in treatment 9, which had more concentration of HP-2MG (a polar resin), the recovery of oxygenated taxanes was significantly increased, reaching a value of 190 ± 20 mg/L (Figure 2B). This higher taxane recovery value was 1.53-fold, 1.84-fold and 1.62-fold higher compared to the use of the single HP-20, XAD7HP and HP-2MG resins, respectively. However, beyond total taxane recovery, it can be seen that the oxygenated ratio in every single treatment was lower compared with the results given in treatment 9 (Figure S10 from the supplementary material). This result is in agreement with previous studies focusing on the P450 for the synthesis of Taxadiene-5α-ol, highlighting that careful tuning is required to optimize the oxygenated ratio of taxanes (Nowrouzi et al., 2022; Nowrouzi & Rios-Solis, 2021).

The change of the concentration in total resin beads also had an important effect for the *in situ* adsorption of total taxanes. For instance, treatments 1, 2, 7 and 13 had the same resin proportion; however, treatment 1 had a total resin concentration of 1.5 % (w/v), which was the lowest total taxane titer from the 4 treatments with 57 ± 12 mg/L. Comparing this treatment with treatment 2, which had the highest resins concentration of 4.5% (w/v), resulted in 148 ± 17 mg/L of total taxanes (increment of 2.6-fold). These results support a previous study that found using HP-20 resins had a detrimental effect on taxadiene recovery for resin concentrations below 3% (w/v) (Santoyo-Garcia et al., 2022). However, not only was the total concentration of resins an important factor for total taxane titers but the proportion of the individual resins was also seen to affect the total taxane yields as well. Treatments 1, 5, 8, 11 and 12, which all possessed a high concentration of HP-20 (1-1.5 % (w/v)) resin and low HP-2MG concentration (0.5 % w/v), showed the lowest taxane recovery. Treatment 11 resulted in the lowest total taxane titer at 39 ± 7 mg/L. On the contrary, the highest titer of total taxanes were found in treatments 2, 3, 4 and 9 where the main constant was the high concentration of HP-2MG resin (1.5 % (w/v)). Treatment 9 had the highest total taxanes titer with 190 ± 20 mg/L, which represented a 4.8–fold increment compared with treatment 11 and 7.9–fold increase compared to the control where no extraction aid was used. Therefore, using 1.5 % (w/v) of total resins with equal resin proportion (treatment 1), resulted in the lowest taxane titer value.

The effect of the three adsorbent resins as factors of the DSD was evaluated by forward stepwise regression following the BIC stopping criterion and no rules. A response surface model was derived, thereby considering all main effects, quadratic and any second-order interactions. The resulting statistical model revealed that HP-2MG resin concentration (w/v) (p < 0.02) had the most significant effect. The statistical model was subsequently used to predict the optimal settings for each of the significant factors, as summarized in Figure 3.

**Figure 3.**
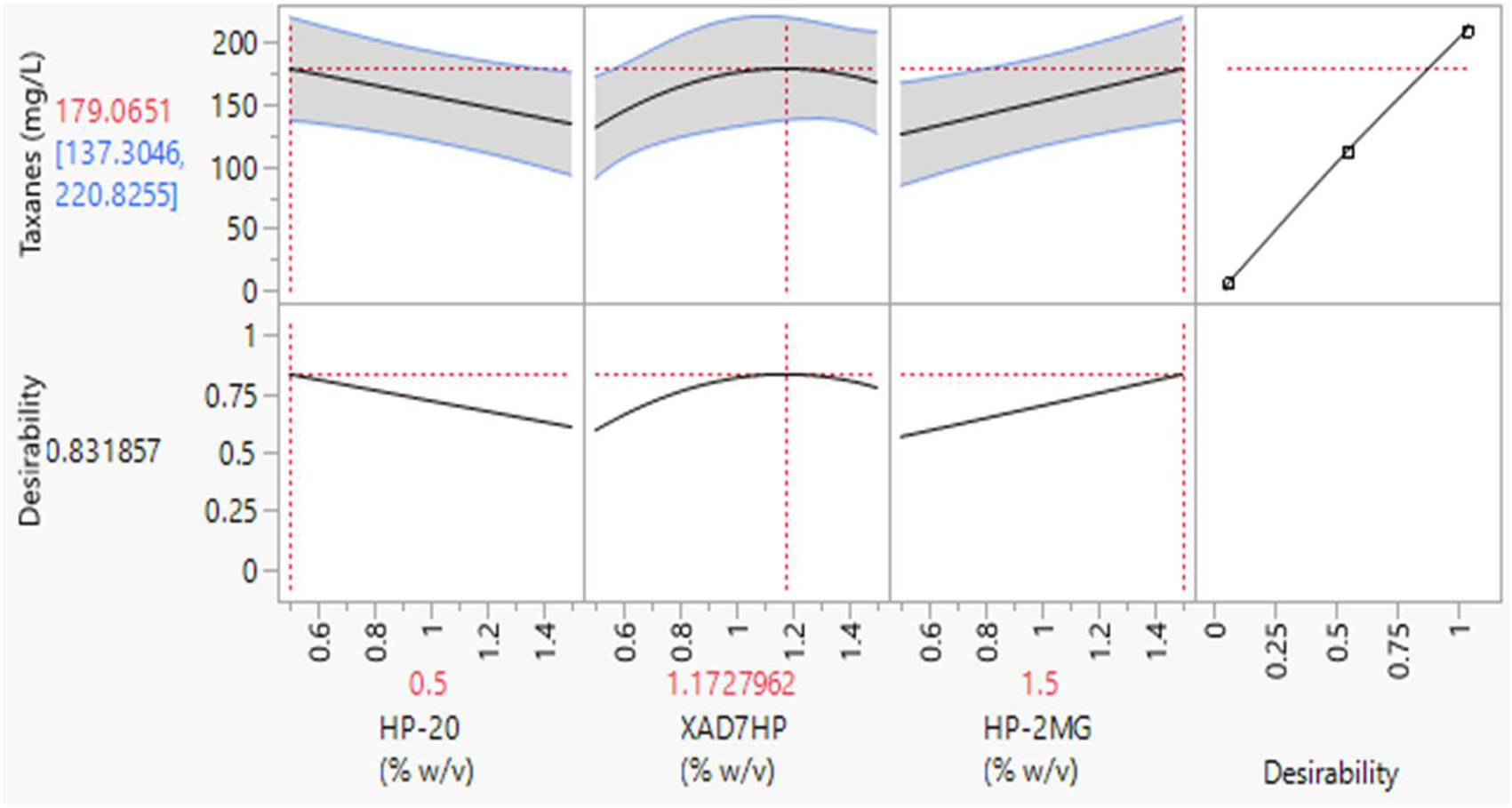
Prediction factors profiler for the optimum resin concentration. The top-half section of the Figure indicates the optimum concentrations of each resin for a maximized desirability (0.83). The first three graphs of the top-half show the optimum resins concentrations (x axis) in red numbers (0.5, 1.172 and 1.5 % w/v for HP-20, XAD7HP and HP-2MG respectively) to obtain the highest taxane concentration, which was predicted to be 179 mg/L (dashed red line in Y axis). The shadowed area between the blue lines of the same graphs indicates the confidence interval for each factor. Graphs in the bottom half indicate the maximum desirability score to maximize taxane recovery.

According to the DoE analysis, the main effect to obtain higher taxane titers was the polar resin HP-2MG. Therefore, the concentration of the HP-2MG resin should be kept at the upper level of the treatments tested, which was 1.5% (w/v) (Figure 3). Interestingly, the model also showed a curved behavior for the aliphatic resin XAD7HP, making the optimum concentration for that resin to be 1.2% (w/v). Lastly, the prediction profiler for the selected model showed that the non-polar resin concentration of HP-20 should be left at the minimum tested concertation of 0.5% (w/v). Therefore, the best resin combination of adsorbent resins in the cultivation to obtain the higher taxanes titers according to the selected model was 0.5, 1.2 and 1.5% (w/v) of the resins HP-20, XAD7HP and HP-2MG, respectively. This result agrees with the combination that was set up in treatment 9, with a small difference in the concentration of the XAD7HP resin which had 1% (w/v) in treatment 9 and 1.2% (w/v) in the optimized model.

To further investigate taxane adsorption using the ISPR, the same protocol for cultivation and extraction was performed, with the exception that the resins were separated from the media and biomass for the final quantification step. Only treatments that gave the best taxane titers were run (Treatments 2, 7 and 9). Results are detailed in Figure 4, where taxanes adsorbed by the resins represented 80% on average of the total taxanes produced after the cultivation period, leaving the rest in the media and biomass.

**Figure 4.**
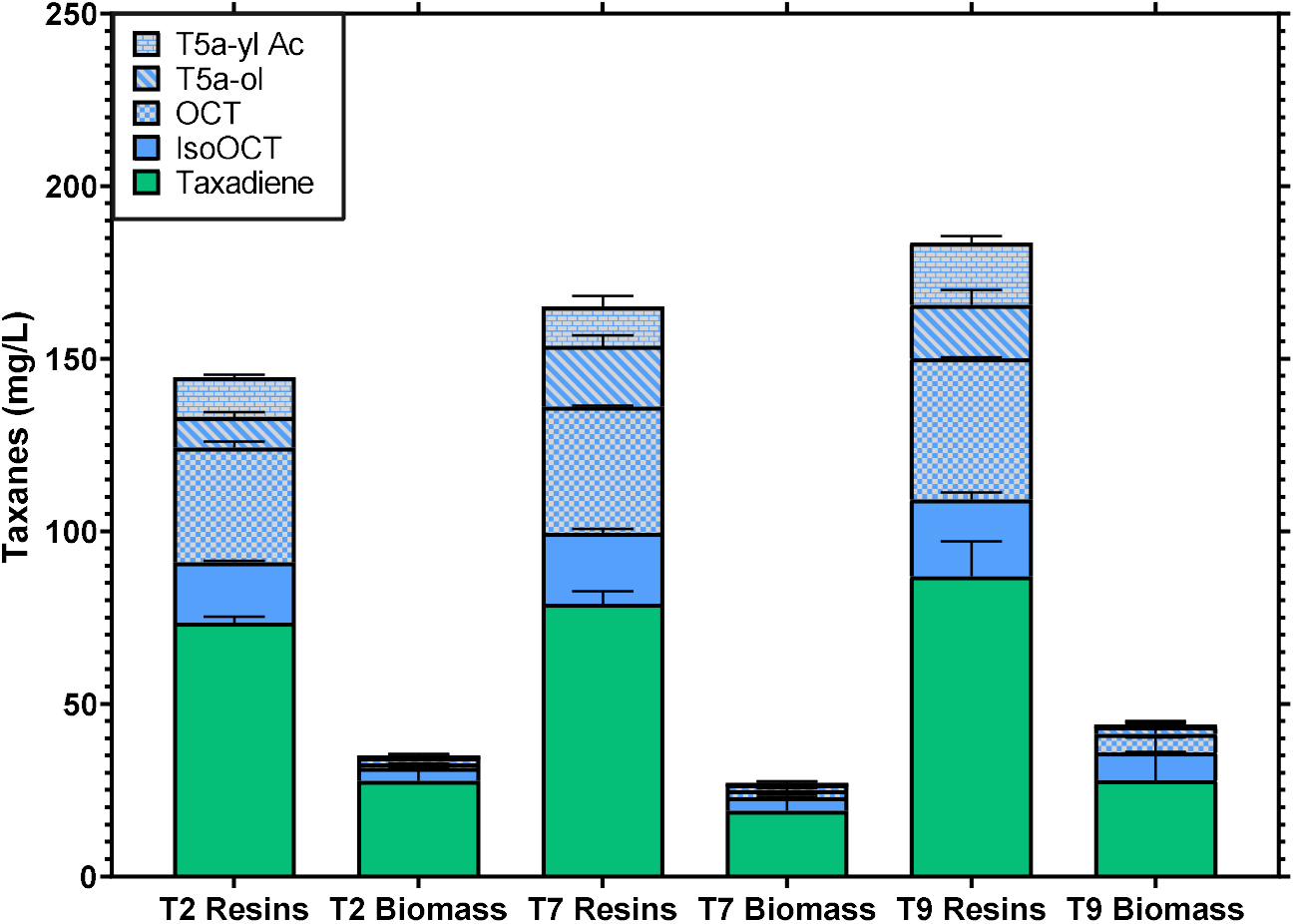
Taxane concentration in the partition between resin and biomass in Treatments 2, 7 and 9. All treatments had the same cultivation parameters of 30 °C, 350 rpm shaking speed, 2 mL of the working volume and media composition, with the only difference being the composition of the three resins in each treatment (Table 1). Both biomass and resins had the same extraction procedure using 99% acetone. Bars represent the mean, and the error bars the standard deviation (n = 2).

Total taxanes were distributed almost equivalently in the three selected treatments, which were retrieved into the resins up to 144 ± 7, 165 ± 18 and 184 ± 21 mg/L, whilst having a total taxane concentration in the biomass up to 35 ± 2, 27 ± 1.5 and 44 ± 5 mg/L from treatments 2, 7 and 9 respectively (Figure 4). These results represent a partition into the resins for 80, 85 and 80% of the total taxanes for treatments 2, 7 and 9 respectively. These results support previous studies where sesquiterpene (+)-zizaene was recovered *in situ* using HP-20 and XAD4 resins (used independently from each other) produced by engineered *E. coli*, where around 86% and 82% of the compound was recovered by the adsorbent material HP-20 and XAD4, respectively (Aguilar et al., 2019). In very similar studies, where only taxadiene was produced by gene-edited yeast and retrieved using only HP-20 resin, the partition was 50% between the resin and biomass (Santoyo-Garcia et al., 2022). This difference from our previous results could indicate that by combining adsorbent materials in the cultivation, the recovery of the product of interest from cells can be improved in comparison to using only a single adsorbent resin (Santoyo-Garcia et al., 2022).

### 3.2 Natural products discovery

An interesting advantage of this methodology is the higher capacity to recover many different natural products or secondary metabolites, since, as mentioned, it has the benefit of retrieving metabolites that had been secreted along with intracellular metabolites, leveraging the extended absorbing capacity of the combined beads. When compared to other retrieval methods (no extraction aid and the most commonly used the dodecane overlay method), the *in situ* cultivation using different resins showcased not only a higher total taxanes titer as previously discussed, but it also retrieved more taxane compounds in the GC-MS chromatograms (Figure 5).

**Figure 5.**
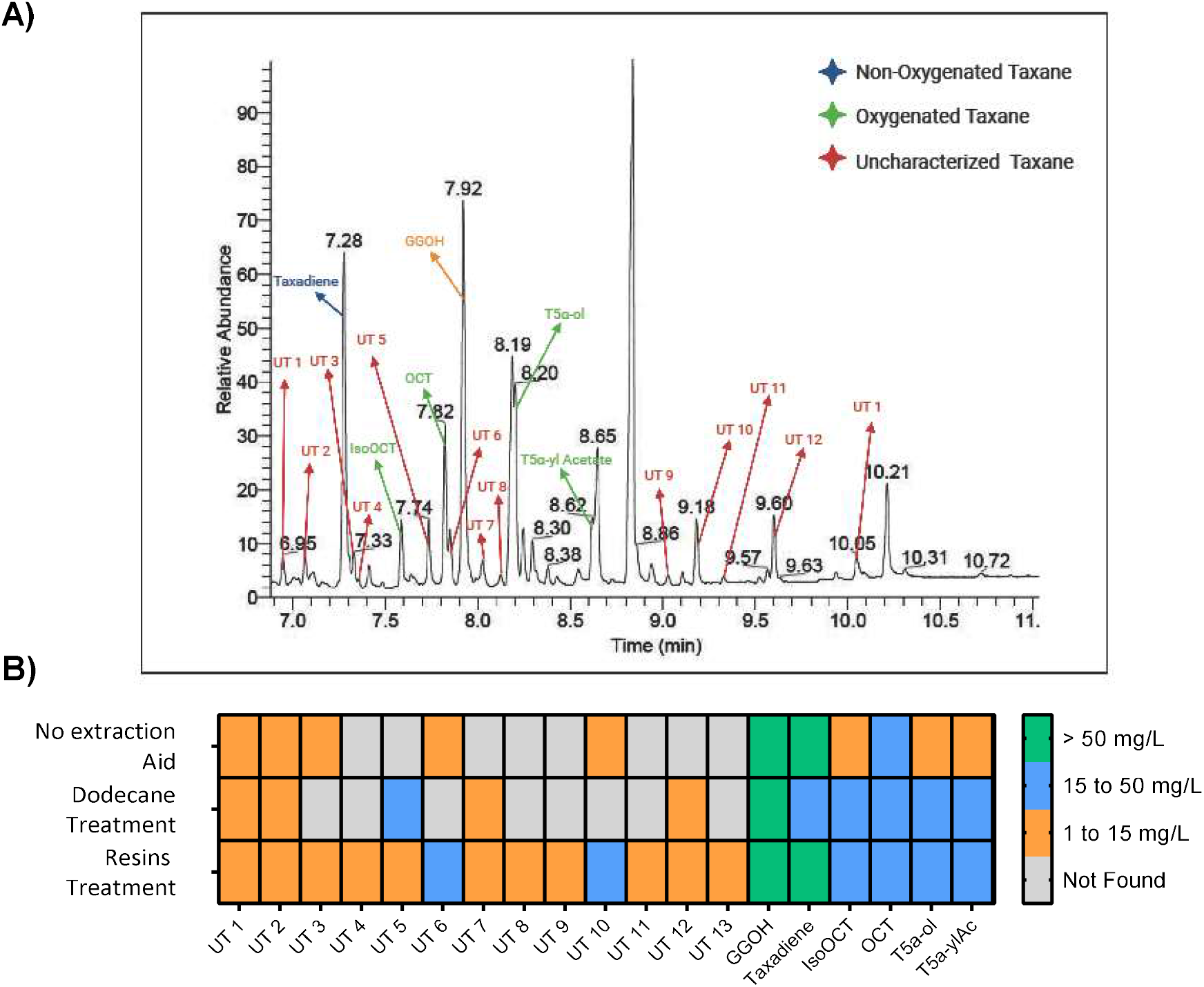
Total taxanes seen in different recovery methods for strain *EJ2*. A. Chromatogram of treatment 9 using the *in situ* cultivation method with resins combination, where the UT labels stand for ‘Uncharacterized Taxane’ for possible new taxanes. B. Heatmap for the taxanes presented in the GC-MS analyses of three methods (no extraction aid, dodecane overlay treatment and *in situ* resins treatment) as well as the concentrations of these taxanes.

In total, 18 taxanes were found in treatment 9 with the resin’s combination of 0.5, 1 and 1.5% (w/v) of HP-20, XAD7HP and HP-2MG respectively (Figure 5A). Using the same cultivation conditions, a dodecane cultivation and a cultivation without any extraction aid were run, from which only 11 and 10 total taxanes were found respectively in their GC-MS chromatograms (Figure S2 of supplementary material). We are reasonably confident that the compounds labelled as ‘uncharacterized taxanes’ could be new taxanes produced by the cell factories, as they had a similar chemical structure to taxanes, showing the key backbone of taxanes’ mass spectra (for instance, abundance in peaks 273, and 288 m/z) (Figures S3 to S6 of supplementary material). The increased presence of taxanes in the combined resin treatment could be explained by the fact that most of the taxanes synthesized by the yeast strain *EJ2* were poorly secreted or not secreted at all to the media. Besides this, the limitations of the dodecane overlay method (such as the evaporation of dodecane during cultivation) could make it harder for GC-MS detection. In previous studies using similar yeast strains for the recovery of taxanes and a dodecane overlay method, only 8 (Walls et al., 2021) and 7 (Nowrouzi et al., 2022) taxane compounds were detected in comparison to the 13 taxane compounds found in treatment 9. This highlights the significant benefits of this method for research focusing on the screening and discovery of new natural products.

### 3.3 Cell-waste removal of ammonia and reactive oxygen species (ROS)

The removal of different cell-waste products using the beads was evaluated in the *S. cerevisiae* strain *EJ2*. The main results observed were a clear reduction of ROS and a slight decrease in ammonia present in the media (Figure 6).

**Figure 6.**
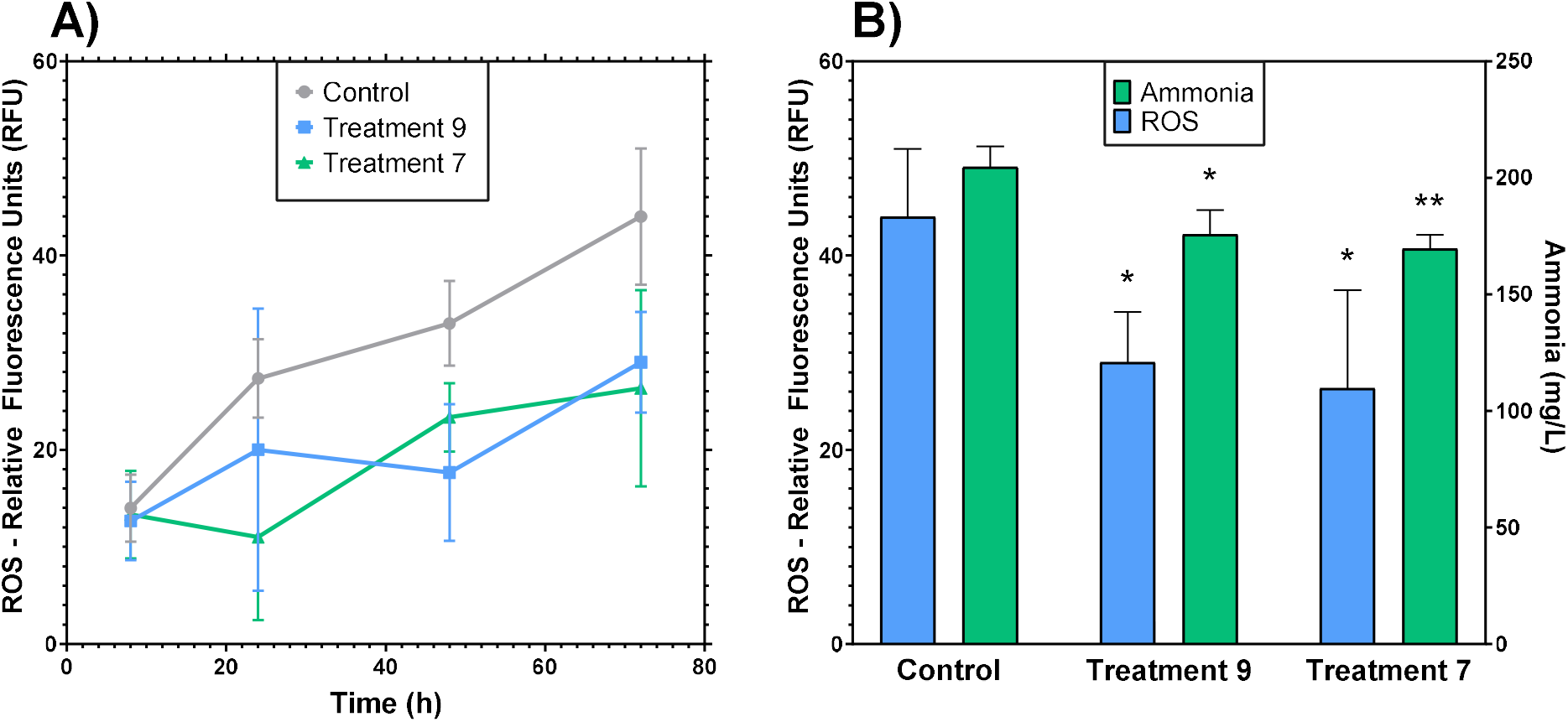
Cell-waste concentrations of ammonia and ROS during cultivation of strain *EJ2*. A) Kinetics of ROS expressed in relative fluorescence units (RFU) of the oxidized compound H_2_-DCFDA during the 72 h cultivations of treatments 9 and 7 (both treatments with a total of 3% (w/v) of total resins), as well as the control with no *in situ* resin. B) Emitted fluorescence of intracellular ROS and ammonia concentrations of the same treatments at the end of each cultivation (72 h). Bar and point values represent the mean (n = 3). Error bars represent the S.D. Statistical significance was assessed by Dunnett’s test versus their corresponding controls. *P <0.0332; **P <0.0021; n.s.= not significant P>0.1234.

The final ammonia concentrations in treatments 9 and 7 were 176 ± 8 and 169 ± 5 mg/L respectively, compared to the control value of 205 ± 7 mg/L at the end of cultivation (Figure 6B). The accumulation of ammonia in the control with no adsorption aid was 1.15-fold higher than treatment 9 and 1.2-fold higher than treatment 7, indicating that the interaction with this chemical during cultivation is reduced. This suggests that cell viability and optimal culture conditions are improved in resin treatments (Figure 6B). In previous studies, it has been demonstrated that ammonia both reduces yeast growth at high concentrations and can inhibit important enzymes, such as Acyl-CoA Synthases (ACS) (Zheng et al., 2012). This could also explain why total taxane production was enhanced by the resin treatments, as acetyl CoA is the initial step for terpenoid synthesis in strain *EJ2*. Increasing ammonia from 337 to 673 mg/L produced an increase of pH in *Cryptococcus c*. yeast from 7.5 to 8.5, having an inhibitory growth effect towards the production of a biofuel (Zheng et al., 2012). For this reason, trying to maintain ammonia levels near the initial cultivation concentration could help optimize cell growth and desired NP production.

In contrast, ROS levels were statistically higher in control cultivations (Figure 6A) compared to treatments with beads after 24 hours of cultivation. After 72 hours, the difference between the fluorescence emitted by the resin combination treatments was on average 30 ± 6 compared with 44 ± 5 in the control, meaning ROS concentration was 1.5-fold lower in treatments 9 and 7. Interestingly, no statistical difference was seen between the two resin combination treatments (Figure 6A). ROS production by yeast has an important effect on its growth and cell viability. *Tert*-butyl hydroperoxide (*t*-BuOOH), hydrogen peroxide (H_2_O_2_) and menadione are common ROS produced by *S. cerevisiae*, and it has been shown that ROS concentrations greater than 0.4 mM or 0.6 mM can inhibit the growth of these yeast (Gazdag et al., 2014). Importantly, engineered *S. cerevisiae* could also induce the production of additional ROS by the action of cytochrome P450s enzymes (CYPs), which are necessary for the synthesis of oxygenated taxanes, provoking a pro-oxidative imbalance (Zhou et al., 2015).

### 3.4 Semi-continuous cultivations at microscale

Using the same resin combination of treatment 9, a set of runs was conducted using a microplate to evaluate the possibility of semi-continuous production of taxanes. Cultivation parameters were not changed in comparison to batch cultivations; the main difference was the removal of spent media and biomass before reaching the stationary phase for new fresh YPG media in each cycle after 48 hrs. In addition, 3, 6, 9 and 12% (w/v) total resin bead concentrations were used, keeping the same bead ratio of treatment 9 (0.5, 1 and 1.5% (w/v) of HP-20, XAD7HP and HP-2MG, respectively). The kinetics of biomass and total taxane production was evaluated throughout 3 cycles of media replacement and the total taxanes were recovered and compared with the batch cultivations (Figure 7).

**Figure 7.**
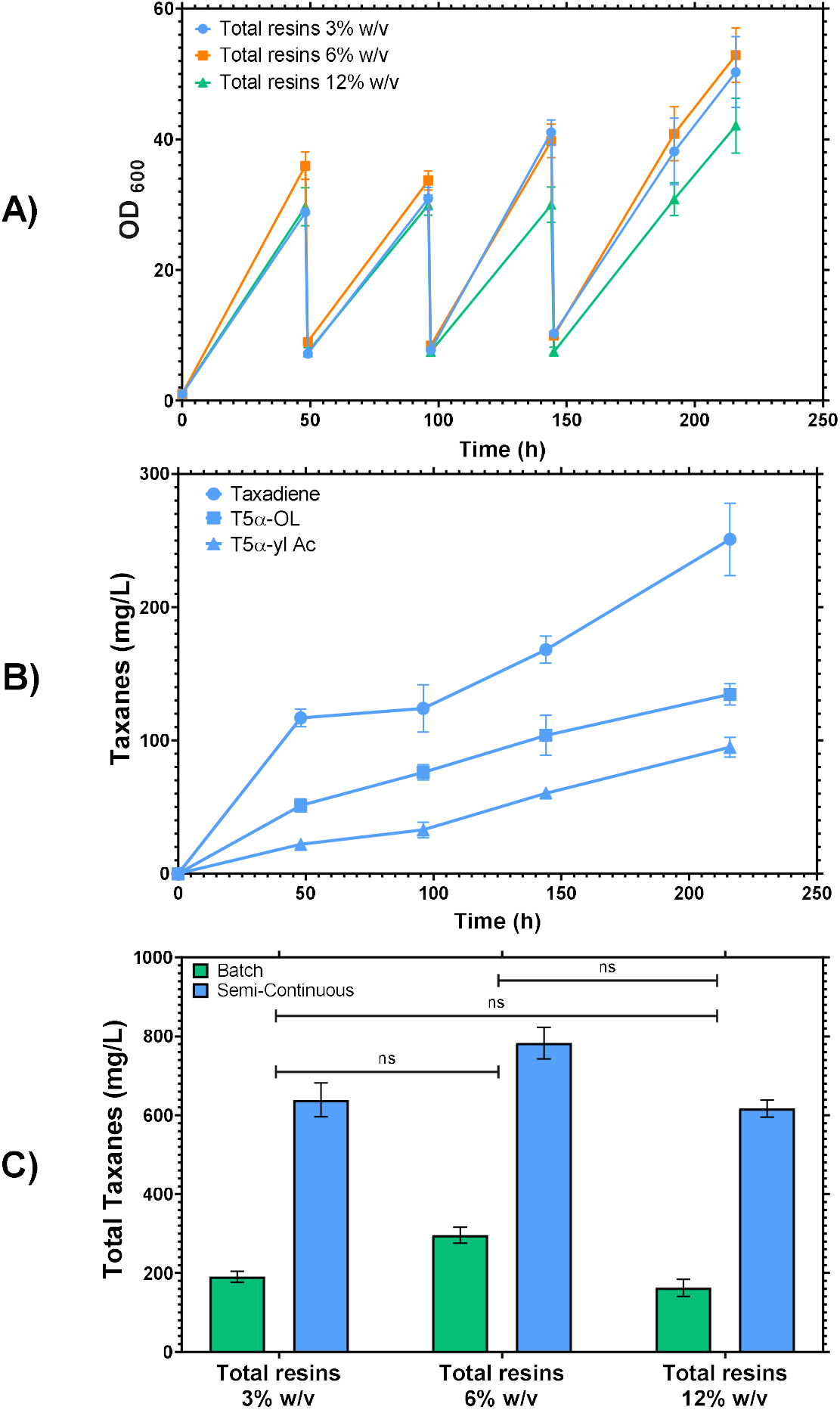
Biomass kinetics and total taxane production for the semi-continuous microplate cultivations. A) Biomass kinetics during the cultivation time for 3 cycles of media replacement. The cultivations had the same parameters as batch cultivations with 30 °C, 350 rpm and media YPG concentrations. B) Kinetic production of the paclitaxel precursor’s taxadiene, T5α-ol and T5α-yl Acetate, for treatment 9 during the semi-continuous cultivation using 6% (w/v) resins. C) Total taxanes recovered from treatment 9 maintaining the same three resin proportions for 3, 6 and 12% (w/v). Green bars represent the total taxane concentrations (mg/L) for batch cultivations and blue bars the total taxane concentrations (mg/L) for semi-continuous cultivations. Bar and kinetic point values represent the mean (n = 3), and error bars represent the S.D.

Before the removal of the first media, biomass was around OD_600_ = 35 and 37 in the experiments, with total resin amounts of 3% and 6% (w/v), respectively, and around OD_600_ = 30 for 12% (w/v). At the end of the semi-continuous cultivations, the final biomasses were OD_600_ = 50 ± 4, 53 ± 3 and 42 ± 3, for 3, 6 and 12% total resins (w/v), respectively (Figure 6A). This biomass behavior at the end of the semi-continuous cultivation was similar to that obtained in the batch cultivations, which indicates that the renewal cycles did not interfere with biomass generation. Another important result seen in Figure 7A is how the engineered strain *EJ2* endured the high resin concentration of the 12% (w/v) treatment. It has been previously reported that a very similar strain (*LRS5*), which could only produce taxadiene due to its heterologous expression, did not grow on a 12% (w/v) cultivation using only HP-20 resin beads in microplate cultivation at the same conditions (Santoyo-Garcia et al., 2022). This result could demonstrate that the combination of resin beads in the cultivation of a delicate strain could be used to diminish the mechanical stress that the *in situ* cultivation might provoke.

Total taxane accumulation was the key result in this approach, as the resins were maintained during the whole semi-continuous cultivation, with the resins managing to continuously adsorb more quantities of taxanes in each cycle. This method highlights the suitability of using microscale tools to characterize fed-batch and semi-continuous fermentation as previously described (Teworte et al., 2022). Another important result was the paclitaxel early taxadiene precursor titers, T5α-ol and T5α-yl acetate, which have a final concentration of 251 ± 25, 135 ± 6 and 95 ± 4 mg/L, respectively (Figure 7B). Higher titers of oxygenated taxanes could be crucial when new metabolic pathways are studied to complete the final paclitaxel pathway in heterologous organisms. These final titers of T5α-ol and T5α-yl acetate are 5.7-fold and 25.7-fold higher than titers of the same compounds in previous studies (Walls et al., 2020).

Figure 7C represents the total taxane titers obtained in the cultivations with 3, 6 and 12% (w/v) of beads (same bead proportions of treatment 9) in batch and semi-continuous cultivations. For the final total taxane titers, there was no statistical significance between these treatments, which indicates that 3% (w/v) is an adequate concentration for a low-cost strategy to create an efficient production system. At the same time, up to 3.8-fold increases in total taxane titers were obtained in semi-continuous mode vs batch cultivations, making this semi-continuous approach suitable to increase titers of total taxanes. These results support previous *in situ* recovery studies, where the semi-continuous approach using 3 cycles and X-5 resin was used to attain a 3.2-fold increase of tanshinone diterpenoid volumetric yields (mg/L) against the batch in *Salvia miltiorrhiza* hairy root cultures (Yan et al., 2005). Besides this, as seen in Figure 7C, the total taxanes retrieved from the cultivations reached 783 ± 33 mg/L after 9 continuous days of cultivation. This product recovery behavior could be compared with previous studies, where the adsorption of natural 2-phenylethanol in an ISPR semi-continuous process had concentrations at the end of each cycle of 10.9, 21.1 and 32.3 g/L, showing a constant increase per cycle in the product yield (Wang et al., 2011).

## 4. Conclusions

In this study, it was shown that by combining three macro-porous resins with different chemical properties in the cultivation of engineered *S. cerevisiae*, it was possible to recover more taxane compounds at higher titers. A combination of the three adsorbent resins with 0.5, 1 and 1.5% (w/v) of HP-20, XAD7HP and HP-2MG, respectively, resulted in a 7.9-fold increase in total taxane recovery compared to the control with no extraction aid. Moreover, by using the semi-continuous approach with 6% (w/v) of total resins, the total taxanes increased to a maximum concentration of 782 mg/L, which represents a 4.1-fold increase compared to batch cultivation. Within this total concentration of taxanes, it was found that the concentration of the oxygenated taxanes T5α-ol and T5α-yl acetate were 135 ± 6 and 95 ± 4 mg/L, respectively, the highest titer values reported to date. Treatments with resin combinations resulted in a lower concentration of cell-waste ROS compared to control with no extraction aid, which could lead to improved cell homeostasis during the cultivation and hence increase taxane production. This work, therefore, highlights a practical and novel ISPR method that leverages different combinations of bead materials for the sustainable and enhanced discovery and recovery of high-value products using microbial cell factories.

## Acknowledgements

The authors would like to thank the technicians at The School of Engineering, University of Edinburgh, UK who helped with the GC-MS analyses Mr Stuart Martin and Mr Mark Lauchlan. Thanks for the support from the IBioE technician Miss Katalin Kis. Thanks to Professor Phil Baran’s Lab at The Scripps Research Institute, San Diego, California for providing the taxadiene standard. Thanks to Phil A. Butin for proofreading this Manuscript.

This work was supported by the Mexican government dependence CONACyT (Mexican National Council for Science and Technology) Scholarship reference CVU: 537962 for JHSG, CVU: 537957 for MVC and CVU: 675492 for NJB. This research was supported by the Engineering and Physical Sciences Research Council (Grant number EP/R513209/1) for LEW and the YLSY Program of the Ministry of National Education of the Republic of Turkey for KM. The Royal Society (Grant Number RSG\R1\180345) and The British Council (Grant Number: 527429894).

## Authors Contributions

Jorge H. Santoyo-Garcia (JHSG) designed and performed the microplate experiments, including most of the data analysis. JHSG wrote the manuscript with input from all authors. Laura E. Walls (LEW) contributed to the design of experiments modelling and data analysis. Marissa Valdivia-Cabrera (MVC) contributed with Reactive oxygen species and ammonia quantification experiments. Koray Malci (KM) and Nestor Jonguitud-Borrego (NJB) contributed with the genome engineering experiments and construction of the yeast cell factory *EJ2*. Leonardo Rios-Solis (LRS) and JHSG conceived the idea for this study. Karen J. Halliday (KJH) and LRS coordinated the study.

## Data availability statement

The data supporting this study’s findings are available from the corresponding author upon reasonable request.

