## Supplementary material for "The synergetic effect from the combination of different adsorption resins in batch and semi-continuous cultivations of *S. cerevisiae* cell factories to produce acetylated Taxanes precursors of the anticancer drug Taxol"

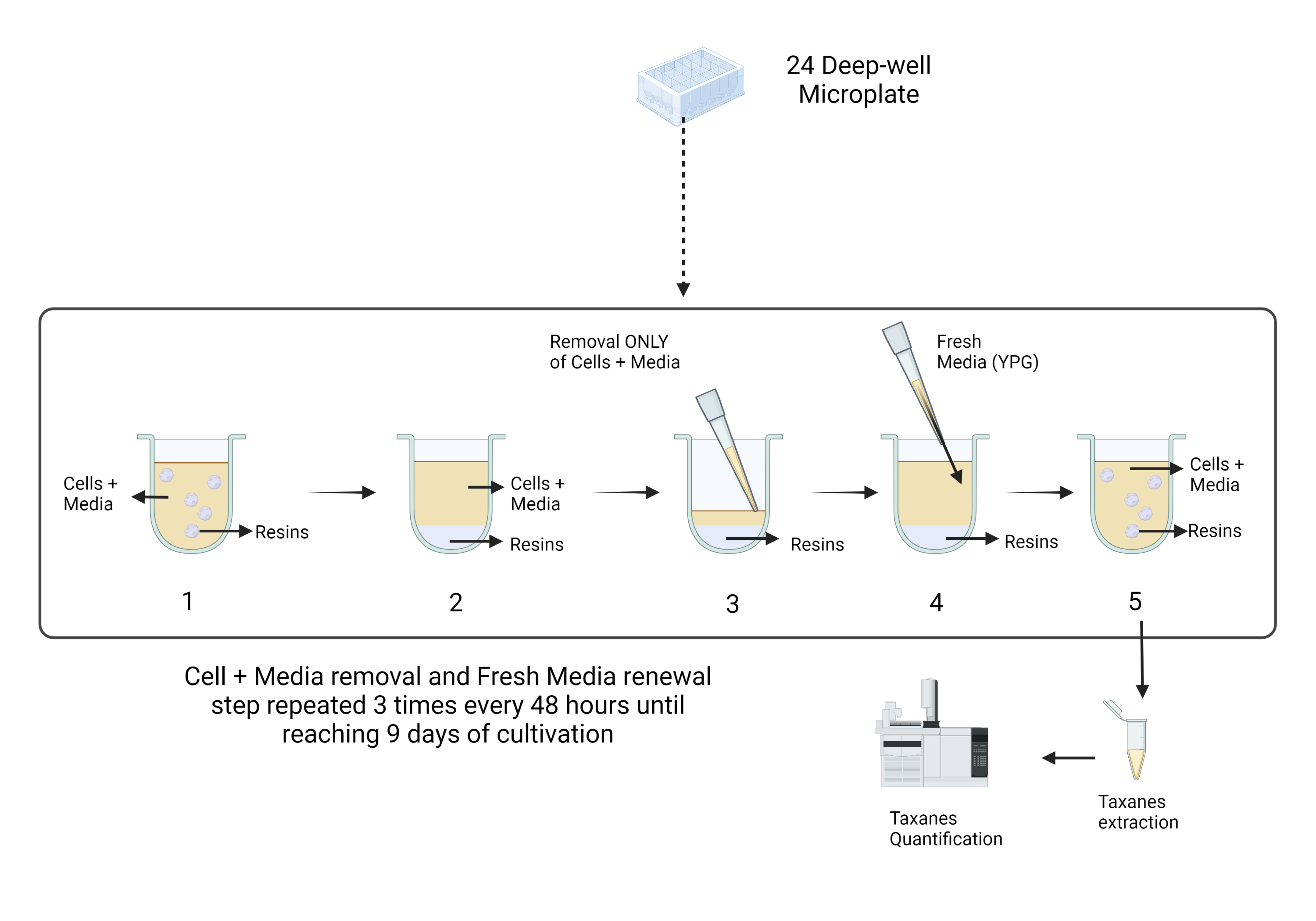

Figure S1. Schematic protocol of Media removal and renewal in the semi-continuous cultivation of *S. cerevisiae* strain EJ2. (1) Happens when each well is mixing in the thermomixer at the cultivation; (2) the microplate is stopped and taken to the bio-hood where after few minutes, the resins will precipitate; (3) removal of 75 % of the cultivation volume without resins is performed (removed volume could be sent to taxanes quantification); (4) the 75 % of the removed volume is refilled with fresh YPG; (5) the microplate is returned to the thermomixer to continue the cultivation until further taxanes quantification.

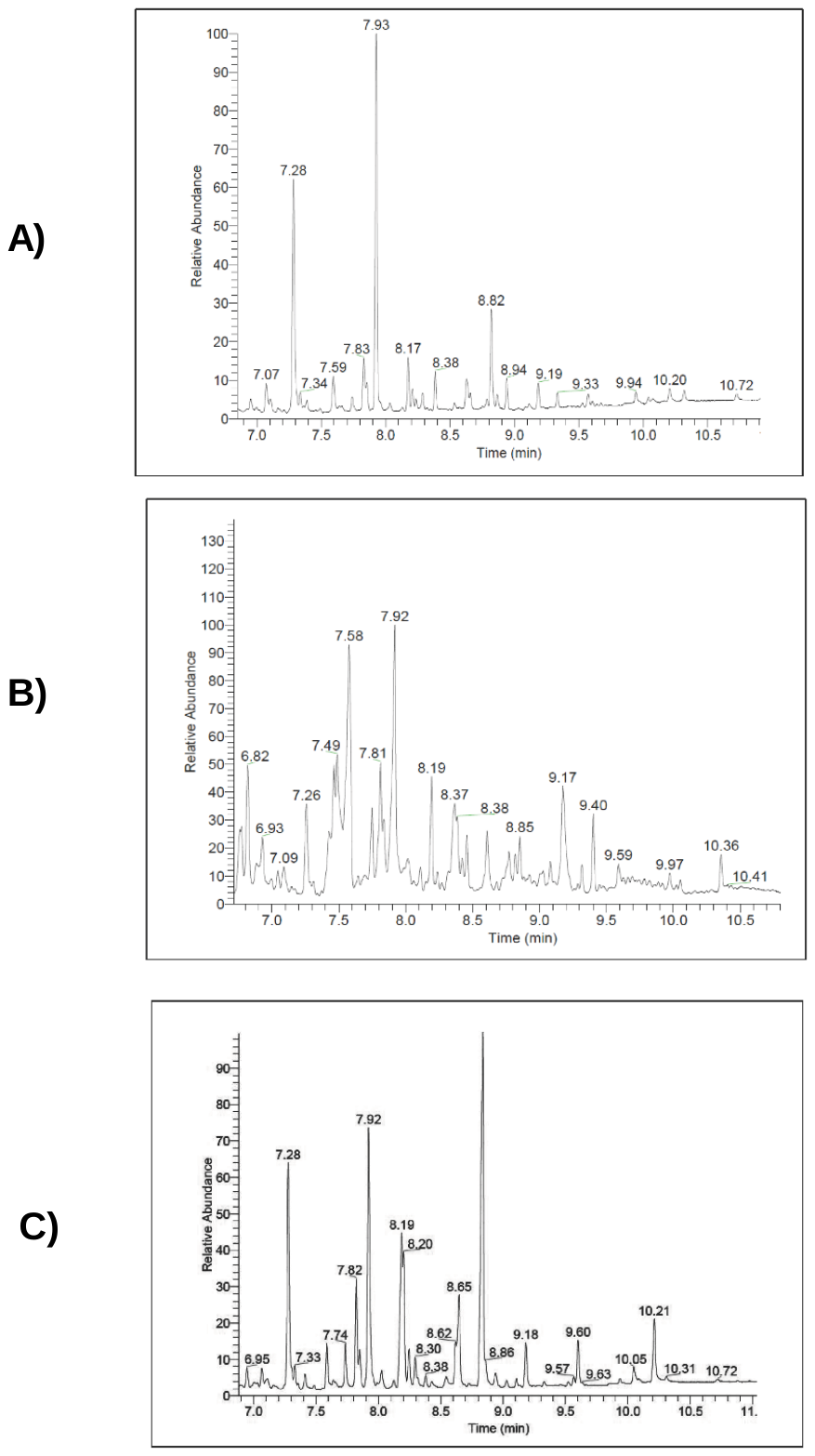

Figure S2. GC chromatograms from different cultivation methods. S2A shows the no extraction aid cultivation with 11 taxanes found. S2B represents the chromatogram form dodecane cultivation where 4 taxanes were fond and S2C showcase the chromatogram for in situ resins cultivation of treatment 9 where 18 taxanes were found.

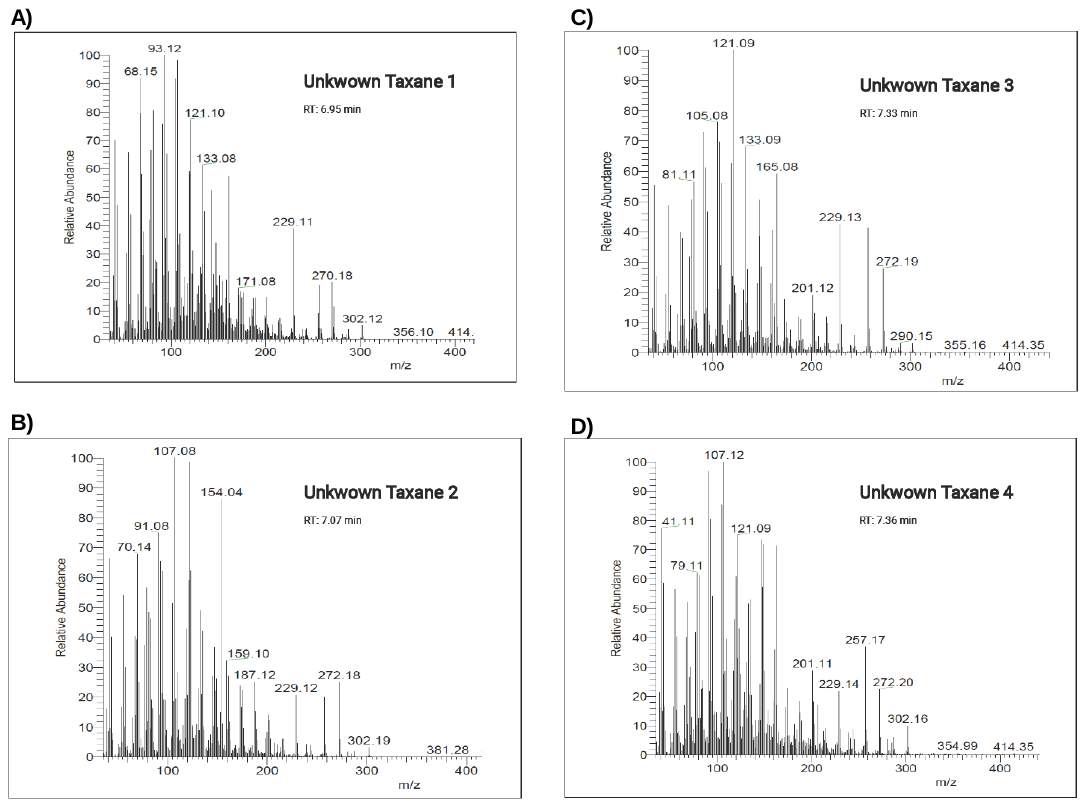

Figure S3. Mass Spectra for unknown taxanes 1 to 4 from treatment 9 of the in situ resins cultivation method. S3A shows unknown taxane 1 with a retention time of 6.95 min. S3B shows unknown taxane 2 with a retention time of 7.07 min. S3C shows unknown taxane 3 with a retention time of 7.33 min. S3D shows unknown taxane 4 with a retention time of 7.36 min.

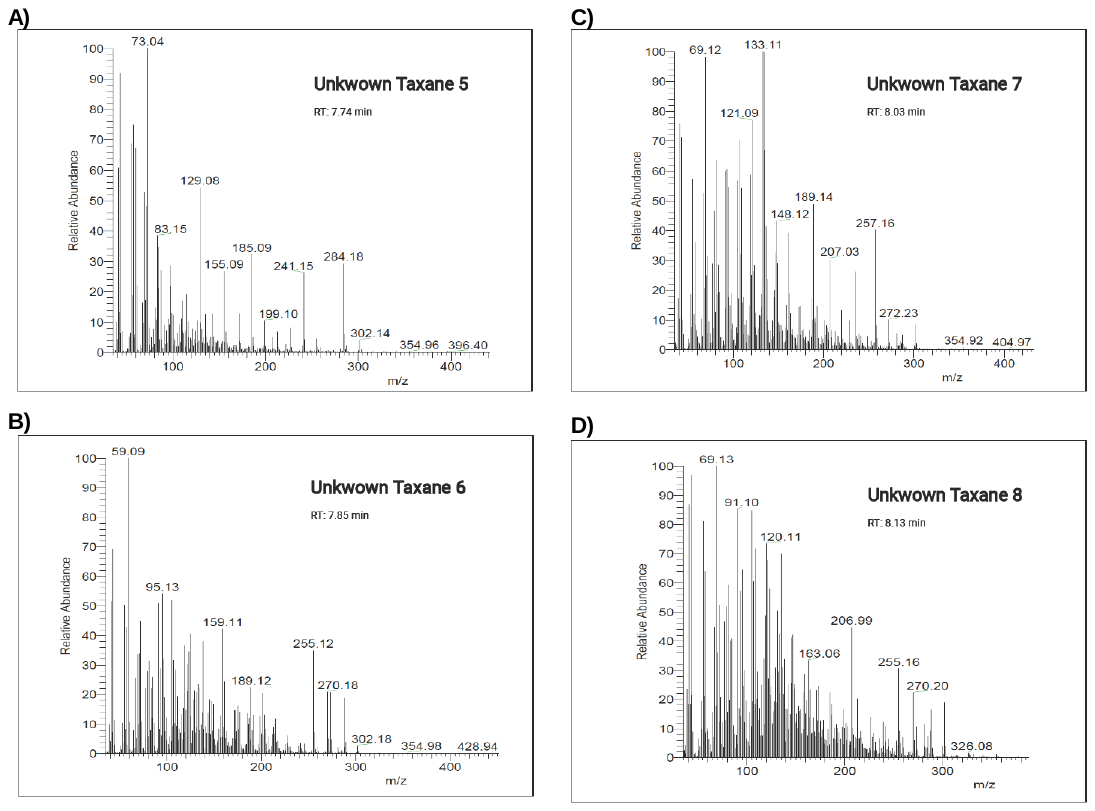

Figure S4. Mass Spectra for unknown taxanes 5 to 8 from treatment 9 of the in situ resins cultivation method. S4A shows unknown taxane 5 with a retention time of 7.74 min. S4B shows unknown taxane 2 with a retention time of 7.85 min. S4C shows unknown taxane 3 with a retention time of 8.03 min. S4D shows unknown taxane 4 with a retention time of 8.33 min.

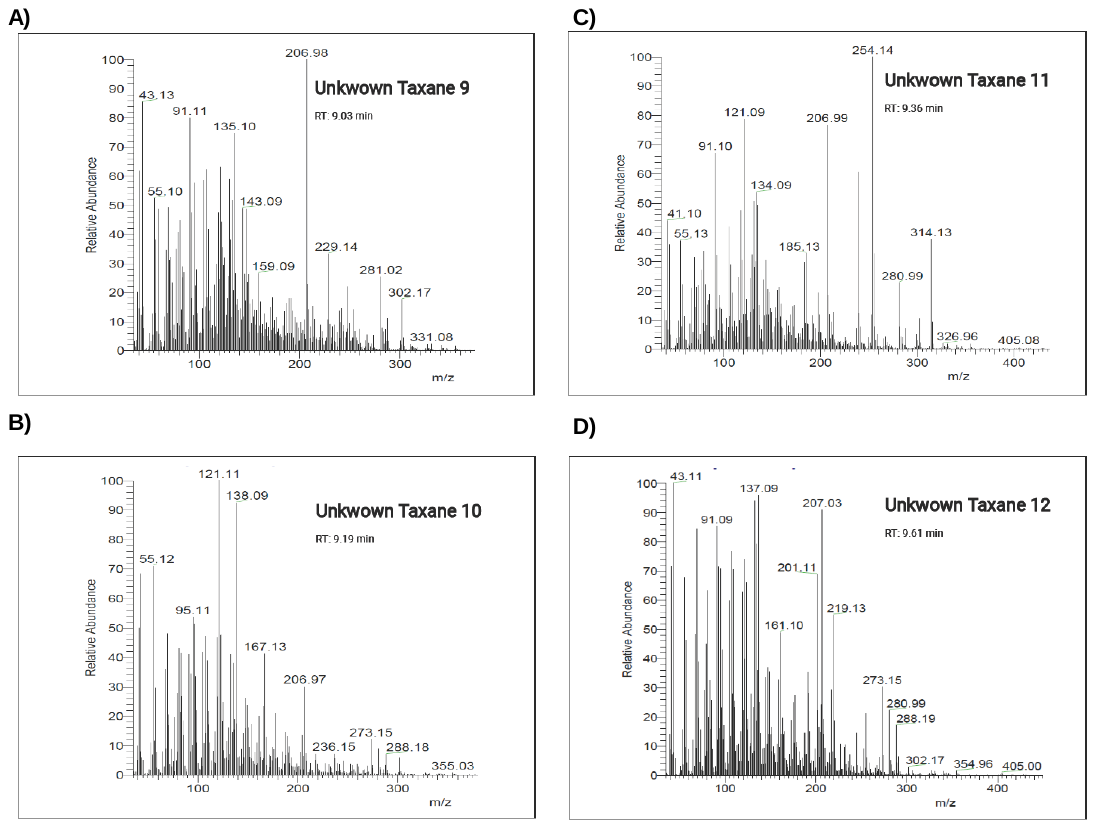

Figure S5. Mass Spectra for unknown taxanes 9 to 12 from treatment 9 of the in situ resins cultivation method. S5A shows unknown taxane 5 with a retention time of 7.74 min. S5B shows unknown taxane 2 with a retention time of 7.85 min. S5C shows unknown taxane 3 with a retention time of 8.03 min. S5D shows unknown taxane 4 with a retention time of 8.33 min.

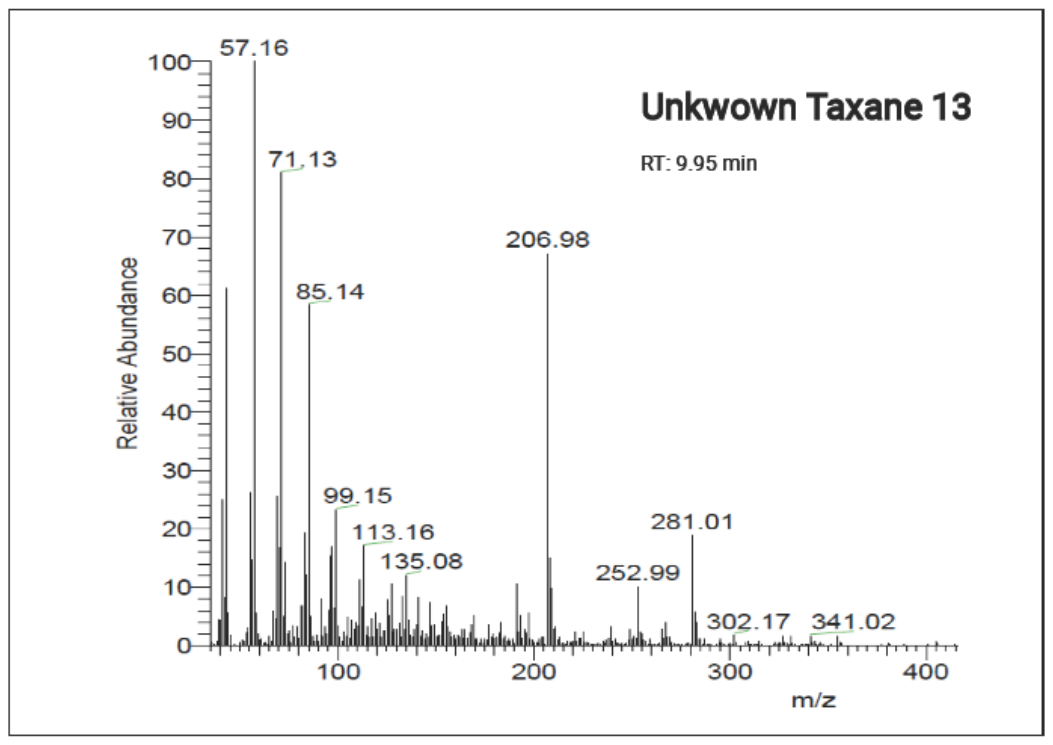

Figure S6. Mass Spectra for unknown taxanes 13 from treatment 9 of the in situ resins cultivation method. Retention time of 9.95 min.

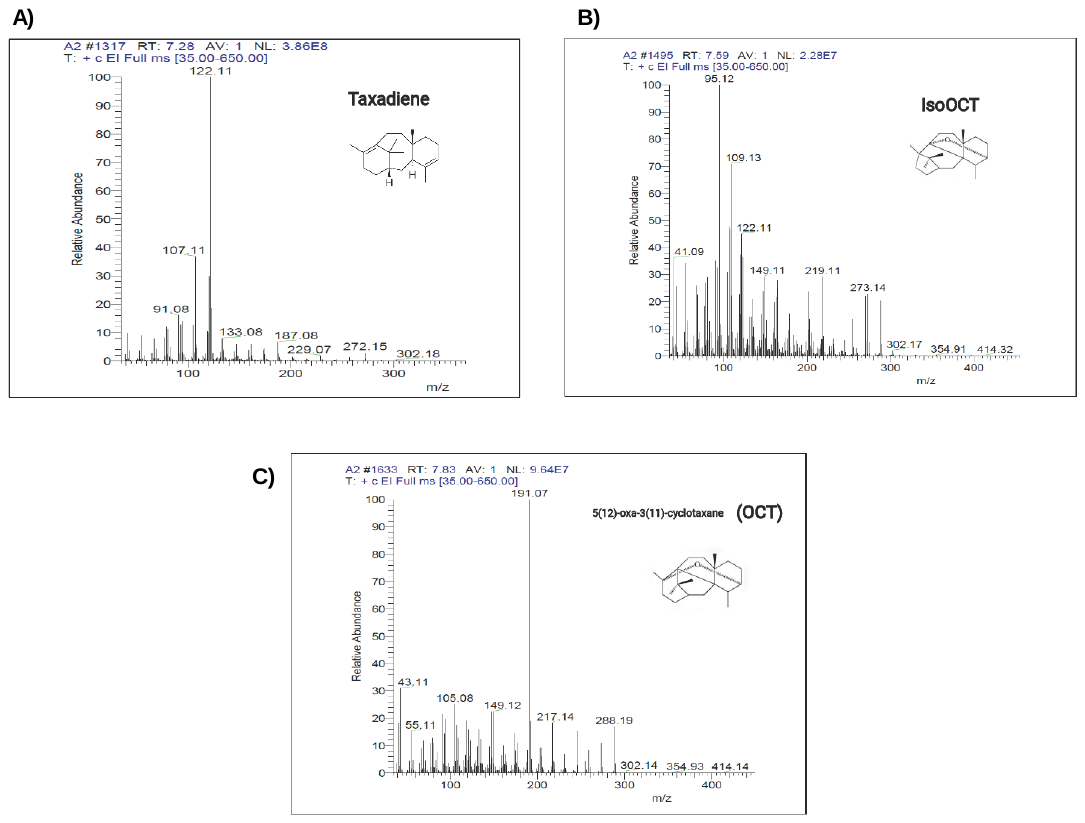

Figure S7. Mass Spectra for S7A Taxadiene, S7B IsoOCT and S7C OCT.

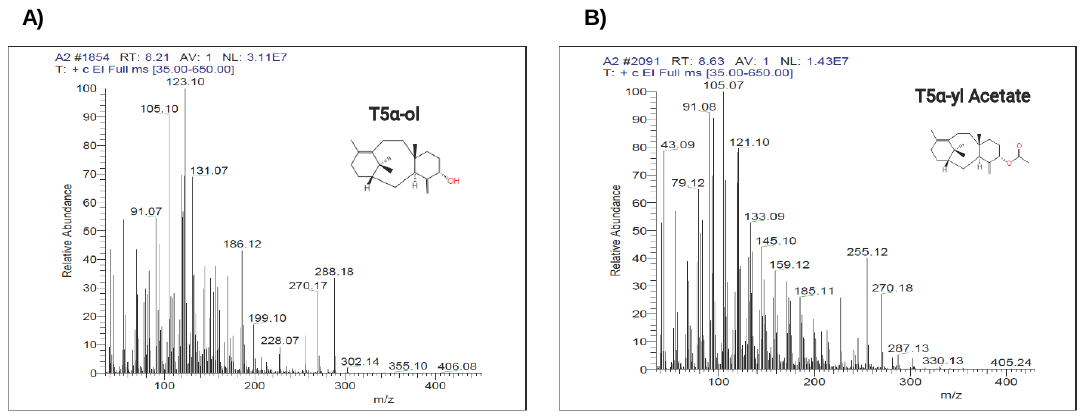

Figure S8. Mass Spectra for S8A T5a-ol and S8B T5a-yl Acetate.

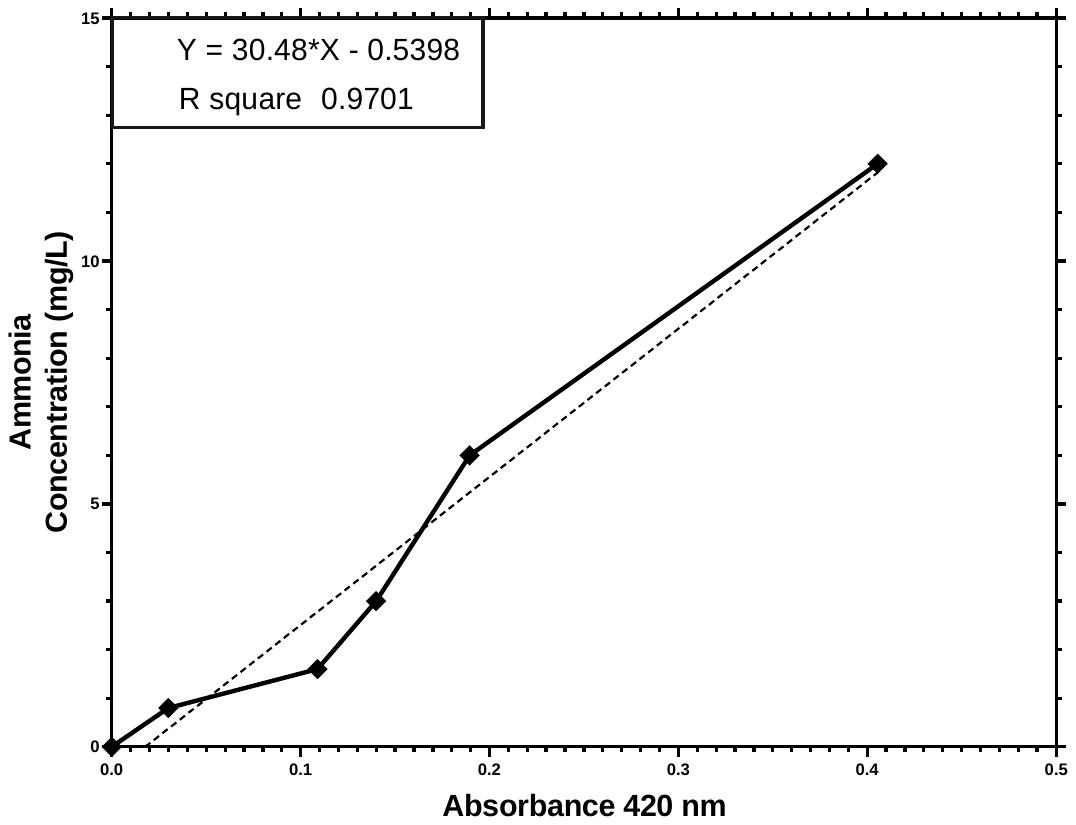

Figure S9. Calibration curve for ammonia quantification using chloride ammonium salts.

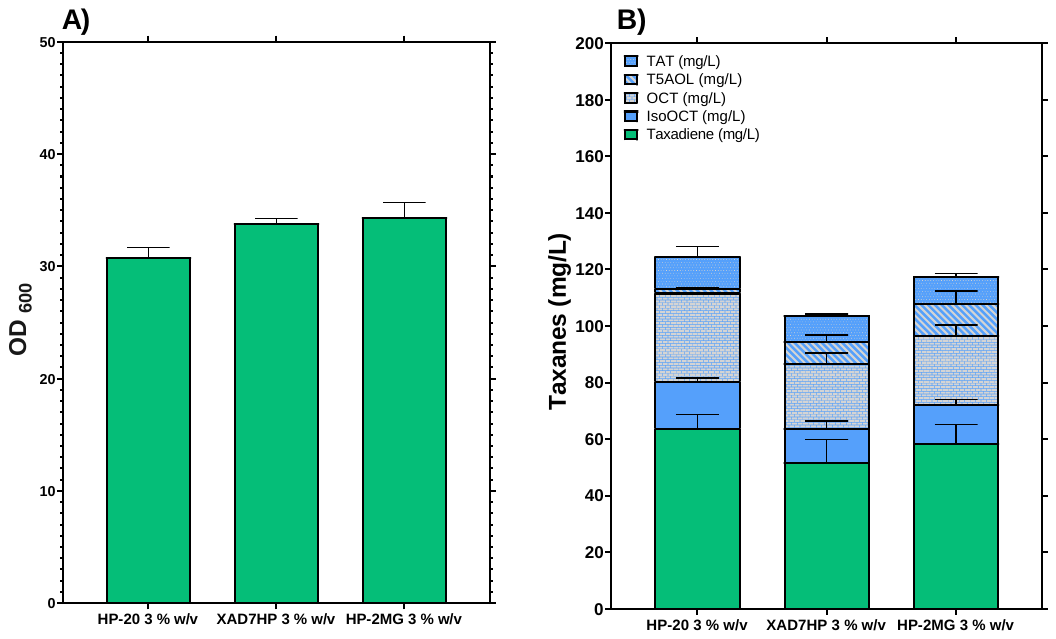

Figure S10. Biomass accumulation at 72 hours of cultivation (A) and Total Taxanes recovered (B) using single resins HP-20, XAD7HP and HP-2MG at 3 % w/v. All treatments had the same cultivation parameters of 30 °C, 350 rpm shaking speed, 2 mL of working volume and media composition. Blue bars represent the oxygenated taxanes and green bars the non-oxygenated taxane. Bar values represent mean ± standard deviation (n = 2).

Table S1. Schematic protocol of semi-continuous cultivation of *S. cerevisiae* strain EJ2. Resin beads are overcharged with produced taxanes in a 10 day cultivation instead of a 3 day cultivation performed in batch cultivation.

| **Days of process** | | | | | | | | | | |
| --- | --- | --- | --- | --- | --- | --- | --- | --- | --- | --- |
| **Cultivation** | | | | | | | | | **Extraction** | |
| 1 | 2 | 3 | 4 | 5 | 6 | 7 | 8 | 9 | 10 | 11 |
| 48 h | | |  |  |  |  |  |  |  |  |
|  |  | 48 h | | |  |  |  |  |  |  |
|  |  | 1^st^  media renewal |  | 48 h | | |  |  |  |  |
|  |  |  |  | 2^nd^ media renewal |  | 72 h | | | | 12 h |
|  |  |  |  |  |  | 3^rd^ media renewal |  |  |  | |
